# Single-trial dynamics of hippocampal spatial representations are modulated by reward value

**DOI:** 10.1101/2020.10.21.349043

**Authors:** Frédéric Michon, Esther Krul, Jyh-Jang Sun, Fabian Kloosterman

## Abstract

Reward value is known to modulate learning speed in spatial memory tasks, but little is known about its influence on the dynamical changes in hippocampal spatial representations. Here, we monitored the trial-to-trial changes in hippocampal place cell activity during the acquisition of place-reward associations with varying reward size. We show a faster reorganization and stabilization of the hippocampal place map when a goal location is associated with a large reward. The reorganization is driven by both rate changes and the appearance and disappearance of place fields. The occurrence of hippocampal replay activity largely followed the dynamics of changes in spatial representations. Replay patterns became more selectively tuned towards behaviorally relevant experiences over the course of learning. These results suggests that high reward value enhances memory retention via accelerating the formation and stabilization of the hippocampal cognitive map and enhancing its reactivation during learning.

## Introduction

Our ability to remember an event depends in part on the behavioral relevance or associated value of the event (Stickgold and Walker 2013). Generally, the more rewarding an experience is, the better and the longer it will be remembered (Salvetti, Morris, and Wang 2014; Gruber et al. 2016; Miendlarzewska, Bavelier, and Schwartz 2016). Rewarding feedback is known to modulate multiple memory processes. In addition to enhancing memory consolidation (Salvetti, Morris, and Wang 2014; Igloi et al. 2015; Gruber et al. 2016; Michon et al. 2019), reward has also been shown to promote memory encoding and to accelerate learning during experience (Wolosin, Zeithamova, and Preston 2012; Igloi et al. 2015; Miendlarzewska, Bavelier, and Schwartz 2016).

The activity of the hippocampus and its interactions with cortical and subcortical brain regions are critical to both the formation and the consolidation of episodic and spatial memory traces (Squire 2004). Brain imaging studies in humans have shown that hippocampal activity and its functional connectivity with other brain regions are modulated by reward magnitude during experience (Wolosin, Zeithamova, and Preston 2012; Igloi et al. 2015; Gruber et al. 2016) and are predictive of future memory recall.

In rodents, during active exploration, individual neurons in the hippocampus are preferentially active when the animal crosses a particular location, or place field, in the environment (O’Keefe and Dostrovsky 1971). Collectively, the activity of place cells form a map-like representation of space. The formation of this cognitive map is experience dependent (Bostock, Muller, and Kubie 1991; Navratilova et al. 2012) and it is sensitive to changes in a large range of spatial and other stimuli, a process referred to as ‘remapping’ (Leutgeb et al. 2004, 2005; Latuske et al. 2018). Notably, studies have reported a number reward-related changes in the firing properties of hippocampal neurons: the accumulation of place fields near newly rewarded locations (Dupret et al. 2010; Danielseon et al. 2016; Tryon et al. 2017), the modulation of place cells firing rate by reward probability expectation (Lee et al. 2012, 2017) and the demonstration of a sub-population of hippocampal neurons that is consistently active at rewarded locations in different environments (Gauthier and Tank 2018). These studies also reported that place cells in the dorsal area CA1 are more sensitive to reward-related changes than place cells in area CA3 (Dupret et al. 2010; Lee et al. 2017). In addition, reward magnitude modulates the reactivation of experience-related hippocampal neural activity patterns (‘replay’) during experience (Singer and Frank 2009; Ambrose, Pfeiffer, and Foster 2016).

However, few studies have investigated the effect of reward magnitude on the hippocampal place code and none has reported an effect (Tabuchi, Mulder, and Wiener 2003; Duvelle et al. 2019), possibly because rewarded locations were kept stationary. Moreover, the impact of reward on the formation and updating of the hippocampal map remains poorly understood. In this study, we take advantage of a recently developed paradigm (Michon et al. 2020) to compare the dynamics of hippocampal code during repeated learning of two rewarded locations associated with different reward size.

## Methods

A total of 6 male Long-Evans rats were used for this study. All experiments were carried out following protocols approved by the KU Leuven animal ethics committee (P119/2015) in accordance with the European Council Directive (2016/63/EU). The results presented here derive from the analysis of a subset of data collected for a previously published study (Michon et al. 2019).

### Behavioral procedure

All rats were trained in a dual reward-place association task (Michon et al. 2020). Rats performed the task on an elevated maze that was split into two environments located at the right and left side of the experimental room. The two environments were separated by a divider and connected to a common home platform via 30 cm long tracks. Each environment consisted of a choice platform with connections for up to 6 radially emanating arms. Each arm ended in a reward platform. The goal in the dual reward-place task is for the animal to learn and remember which of the 6 locations in each environment is associated with either large (9 pellets) or small (1 pellet) reward.

The dual reward-place association task is a repeated acquisition task in which rats need to learn and remember different associations every day. Each daily session is subdvided in an instruction phase and a test phase that were separated by a 2h delay. During the instruction phase, only the rewarded target arm is physically present in each environment. Across 5 trial blocks, rats were allowed to sequentially visit the target arm in the two environments to collect and consume reward. The rats were next placed in an enclosure located in the experimental room for the 2h delay. Following the delay, the rats were tested for their memory of the daily reward-location association separately for both environments in the presence of three additional distractor arms. In each session, task parameters were varied pseudo-randomly, i.e. the location of the target and distractor arms, the large/small reward assignment to left/right environment and the order in which environments were presented to the animal during instruction and test phase.

### Electrophysiological recordings

A custom-designed 3D-printed micro-drive array (Kloosterman et al. 2009; Nguyen et al. 2009), carrying up to 24 tetrodes and 3 stimulation electrodes was surgically attached to the rat skull. During surgery, the array was positioned above the cortical surface through craniotomies located above the dorsal hippocampus for the recording electrodes (center coordinates: 4 mm posterior to Bregma, 2.5 mm right from the midline) and above the ventral hippocampal commissure for the stimulation electrodes (center coordinates: 1.3 mm posterior to Bregma, 0.9 mm right from the midline). Note that the stimulation electrodes were not used as part of this study. Following 1 week of post-operative recovery, the electrodes were lowered towards the pyramidal cell layer of hippocampal area CA1 over the course of 2-3 weeks.

During experiment, electrophysiological recordings were performed using a 128-channel data acquisition system (Digilynx SX, HS-36 analog headstage and Cheetah software; Neuralynx, Bozeman, MO). Wide-band (0.1-6000 Hz) signals and waveform snippets of online detected spikes in the band-pass filtered signal (600-6000 Hz) were sampled at 32 kHz. The position of the rats in the maze was tracked and captured at 50 Hz using an overhead video camera and colored LEDs mounted on the headstage.

### Data Analysis

Analysis of neural and behavioral data was performed using Python and its scientific extension modules (Millman and Aivazis 2011), augmented with custom Python and C++ toolboxes. In particular, we used numpy (Harris et al. 2020) for numerical computations, scipy (Virtanen et al. 2020), pandas (McKinney 2010), statsmodels, scikit-learn and scikit-posthocs for statistical analyses, matplotlib (Hunter 2007) and seaborn for visualization, and Jupyter/IPython (Perez and Granger 2007) for interactive analyses.

#### Behavior

The position of the rats was tracked using an overhead video camera. Running speed was computed as the magnitude of the gaussian (bandwidth 0.5 seconds) smoothed gradient vector of position. In the instruction trials, the average running speed to (from) the reward platforms was computed over the full journey between leaving home (reward platform) and arriving at the reward platform (home).

#### Detection of sharp-wave ripple (SWR) events

The local field potentials from 1-3 tetrodes were downsampled from 32 kHz to 4 kHz and filtered in the ripple frequency band (140-225 Hz). The ripple envelope was computed as the absolute value of the Hilbert-transformed filtered ripple signal, averaged across the selected tetrodes and smoothed with a Gaussian kernel (bandwidth 15 ms). Slow trends in the ripple envelope were removed using a moving median filter (window length 3 seconds). Finally, start and end times of ripple events were detected when the detrended ripple envelope exceeded a low threshold of *μ* + 0.5 × *σ* and the maximum envelope exceeded a high threshold of *μ* + 8 × *σ*. Here, *μ* and *σ* represent the mean and standard deviation of the detrended ripple envelope. Ripple events that were separated by less than 20 ms were merged into a single event, and events with a duration shorter than 40 ms were excluded.

#### Place cell analysis

Spike waveforms were automatically clustered into putative single units using Kilosort2 (https://github.com/MouseLand/Kilosort and manually curated (using software package Phy, https://github.com/cortex-lab/phy) to identify well-isolated and stable single unit activity clusters. Units with a waveform peak-to-trough duration less than 0.5 ms or more than 1% contamination in a 2 ms refractory period were excluded from analysis.

For each unit, we computed the spatial tuning curve as the average firing rate in 1 cm bins along the trajectories from home to reward platform (outbound) and back (inbound). Only time windows in which the run speed exceeded 5 cm/s and no ripple events occurred were included to compute the tuning curves. Separate tuning curves were computed for the trajectories to/from the left and right environments. All tuning curves were smoothed with a Gaussian kernel (bandwidth 10 cm).

Based on the spatial tuning curves, place fields were defined as contiguous locations where the firing rate exceeded a low threshold of 0.1 Hz and the peak firing rate exceeded a threshold of 1 Hz. We used this relatively low threshold of 1 Hz to make sure to include fields that appear or disappear over the five instruction trials. Instead, we set a minimum in-field firing rate of 5 Hz in at least one of the five trials. Finally, narrow place fields (less than 10 cm distance between the outer most spikes in the field) were excluded from the analysis.

#### Speed-corrected in-field firing rates

To remove the relation between run speed and place cell firing rates, we computed a speed-corrected in-field firing rate per instruction trial. First, we model the rate-speed relation by performing a linear regression on all fields across all animals and sessions. In this model, we only included trials and fields with non-zero rates. Next, we computed the corrected in-field firing rate as:

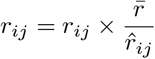

where *r*_*ij*_ is the rate for field *i* in trial *j*, 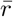 is the mean rate across all fields and trials and 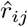 is the speed-predicted rate for field *i* and trial *j*. We used a multiplicative correction to make sure that the corrected rates remained positive. After correction, no linear relation between speed and in-field firing rate remained (supplemental figure S1).

#### Population vector correlation

For each instruction trial, a vector of speed-corrected in-field firing rates was constructed for all place fields or subset of place fields. Trial-to-trial similarity of the population activity was calculated as the Spearman correlation coefficient between the corresponding in-field rate vectors. The population vector correlation was computed at a per-session level and subsequently averaged across sessions.

#### Bayesian neural decoding

Per recording session, an encoding model that relates hippocampal spiking activity to the animal’s position was constructed from the data acquired in the instruction phase of the task. Only spikes emitted during run epochs (run speed > 10 cm/s) with a minimum spike amplitude of 60 *μ*V were incorporated into the model. Tetrodes with a mean spiking rate during run epoch below 0.1 Hz were excluded.

We used a decoding approach that directly relates spike amplitude features to position without prior spike sorting (Kloosterman et al. 2014). Under the assumption that all spikes on a tetrode occur conditionally independent of past spikes and that the firing rate is determined by position in the maze, the hippocampal activity on a single tetrode can be modeled as a marked temporal Poisson process that is fully characterized by the rate function *λ*(**a**, *x*), where **a** represents the vector of spike amplitudes and *x* represent position in the maze. The likelihood of observing a set of spikes with an amplitude of **a**_1:*n*_ in time interval Δ for a given position is then expressed as (Kloosterman et al. 2014):

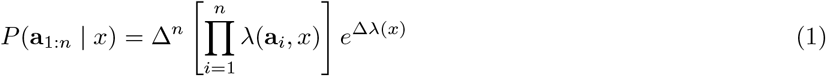

And the joint likelihood across *K* tetrodes is obtained by product of the single tetrode likelihoods: 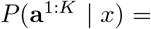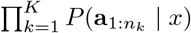

We used a compressed kernel density estimator to evaluate the rate function *λ*(**a***, x*) and the marginal rate function *λ*(*x*) from their component spike count and position occupancy probability distributions (Sodkomkham et al. 2016). The bandwidth of the Gaussian kernel was set to 30 *μ*V for spike amplitude and 5 cm for position. Mahalanobis distance threshold for compression was set to 1.0 for an acceptable trade-off between decoding accuracy and computation time.

We used two different encoding models to decode spatial information in hippocampal replay events. The trial-average encoding model is a single model that incoporates the place field activity from all instruction trials. The single-trial model is composed of a separate model for each of the trial blocks which incorporates only the place field activity within that trial block. For each candidate replay event in a trial block, decoding of the spatial information is performed using the corresponding trial block encoding model.

To perform neural decoding for the spiking activity recorded on *K* tetrodes in time window Δ and estimate the posterior probability distribution over position (sampled at a regular grid with 4 cm spacing) from the likelihood we resort to Bayes’ rule:

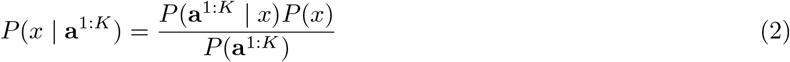

where a uniform prior *P*(*x*) is used.

To evaluate the performance of the decoder, we used a five-fold cross-validation procedure in which four out of five instruction trial blocks were used to build the encoding model and the remaining instruction trial block was used for decoding the animal’s position on the maze in Δ=100 ms time bins. The decoding error was defined as the distance along the track between the estimated and real position. For each session, we evaluated the decoding error distribution separately for left and right environments, as well as for the complete maze. Cross-validation was only performed with the trial-average encoding model. Only sessions with a good decoding performance during the instruction phase (75 percentile of decoding error distribution is below 30 cm) were selected for subsequent analysis of hippocampal replay.

#### Replay analysis

A smoothed multi-unit activity (mua) rate histogram (5 ms bin size, Gaussian kernel with 15 ms bandwidth) was computed from all unsorted spikes recorded from the hippocampus with a peak amplitude larger than 60 *μ*A. The rate was detrended using a moving median filter (window length 3 s). Transient bursts in the detrended mua were defined using a double threshold procedure, where the upper threshold *μ* + 4 × *σ* determines if a burst occurred and the lower threshold *μ* + 0.5 × *σ* determines burst start and end time. Bursts that were separated by less than 20 ms were merged, and bursts with a duration shorter than 80 ms were excluded. Candidate replay events were defined as mua bursts that overlapped with a ripple event and which occurred while the animal was immobile (run speed < 5 cm/s).

Candidate replay events were split into Δ=10 ms time bins and spiking activity (amplitude > 60 *μ*V) in each bin was used to perform decoding as described above. Next, separately for left and right target arms, weighted isotonic regression was performed on the maximum-a-posteriori (MAP) position estimates with posterior probabilities as weights. A goodness-of-fit score was defined that combined both the posterior probabilities and the R^2^ of the regression: 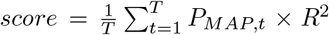, where *T* is the number of time bins in the event, and *P*_*MAP,t*_ is the posterior probability associated with the MAP estimate in time bin *t*. The regression was performed twice to fit both a monotonically increasing and a monotonically decreasing trajectory to the MAP estimates, and only the best fitting trajectory with the highest score was retained.

For each event, the goodness-of-fit score is compared to the distribution of scores constructed from 500 pseudo-random events in which each posterior was randomly drawn from the complete set of candidate replay events (Davidson, Kloosterman, and Wilson 2009). Only candidate replay events with a Monte-Carlo p-value < 0.05 were considered to contain significant trajectory replay.

For each significant replay trajectory, the run direction was decoded using the same Bayesian neural decoding approach as used for position. For each replay event, a direction bias was computed separately for the small and large reward environment as the mean difference in posterior probability between the inbound and outbound run directions across time bins. For each event, the direction bias was compared to the distributions of biases computed from 500 pseudo-random events obtained by randomly assigning each posterior probability of that same event to either the large or small reward environment. Only candidate replay events with a Monte-Carlo p-value < 0.05 were considered to be significantly biased for one run direction. Replay events were classified as either ‘forward’ or ‘reverse’, depending on whether the decoded direction matched the direction of the replay trajectory.

### Statistics

To compare the distributions of trial-to-trial firing rate changes, we used the two-sample Kolmogorov-Smirnov test for equal distributionswith Holm-Sidakp-value correction.

To compare per-session correlation coefficients between trial blocks, we first used the Friedman X^2^ omnibus test, followed by a Conover posthoc test of all pairwise combinations that uses Holm-Sidak p-value correction method.

To compare the number of replay events between successive pairs of trial blocks and between reward conditions, we used the Wilcoxon signed-rank test with Holm-Sidak p-value correction to account for the multiple tests.

To compare the contribution of place cells to SWRs and replay events between pairs of successive trials blocks and between rewards conditions, we used the Mann-Whitney U test followed by a Holm-Sidak p-value correction to account for the multiple tests.

To test the distribution of trial blocks at which individual place fields appear or disappear, we performed a X^2^ test for the null hypothesis of a uniform distribution.

We use p* to indicate p-values that were corrected for multiple tests.

#### Curve fitting

The relation between trial-to-trial correlation or |Δ_*rate*_| and the sequence number of the trial block pair was fitted with a sigmoid growth function:

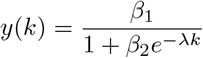

where, *β*_1_, *β*_2_ and *λ* are the parameters to be fitted, and *k ∈* [0, 1, 2, 3] is the sequence number for the trial block pairs (i.e. *k* = 0 for *r*_1→2_, *k* = 1 for *r*_2→3_, etc.). Fitting was performed using the mean squared error and L2 regularization as the cost function. Ten-fold cross-validation was used to estimate the hyperparameter for the L2 regularization. The 95% confidence intervals of the fitted parameters were computed by bootstrapping (1500 samples).

To compare the *λ* parameter between small and large reward conditions, the 95% bootstrapped confidence interval of the difference *λ*_small_ − *λ*_*large*_ was computed. The difference was determined to be significant if the confidence interval did not overlap with zero.

## Results

### Rapid reorganization of spatial representations

A total of 6 rats were trained on a dual reward-place association task (Michon et al. 2019, 2020) in which they learned across 5 instruction trials that two locations were associated with small or large reward (**figure 1a**). The run speed profile across the 5 instruction trials indicated that the rats quickly distinguished the small and large reward destinations (**figure 1b**). On outbound journeys towards the large and small reward, the average run speed steeply increased from trial block 1 → 2. (**figure 1b, left**; Wilcoxon signed-rank test for equal run speed in trial blocks 1 and 2, small: W=27.00, p*=1.5×10^−5^; large: W=18.00, p*=7.1×10^−6^). However, while the average speed towards the large reward remained stable, it progressively decreased for the small reward condition such that the average run speed towards the large reward was significantly higher from trial block 3 onward (Wilcoxon signed-rank test for equal run speed in small and large conditions: trial block 3: W=114.00, p*=0.0051; trial block 4: W=27.00, p*=1.9×10^−5^; trial block 5: W=30.00, p*=1.9×10^−5^). The average run speed on inbound journeys back from the rewarded locations to home mostly remained stable across trial blocks and was not different between reward conditions (**figure 1b,right**).

**Figure 1:**
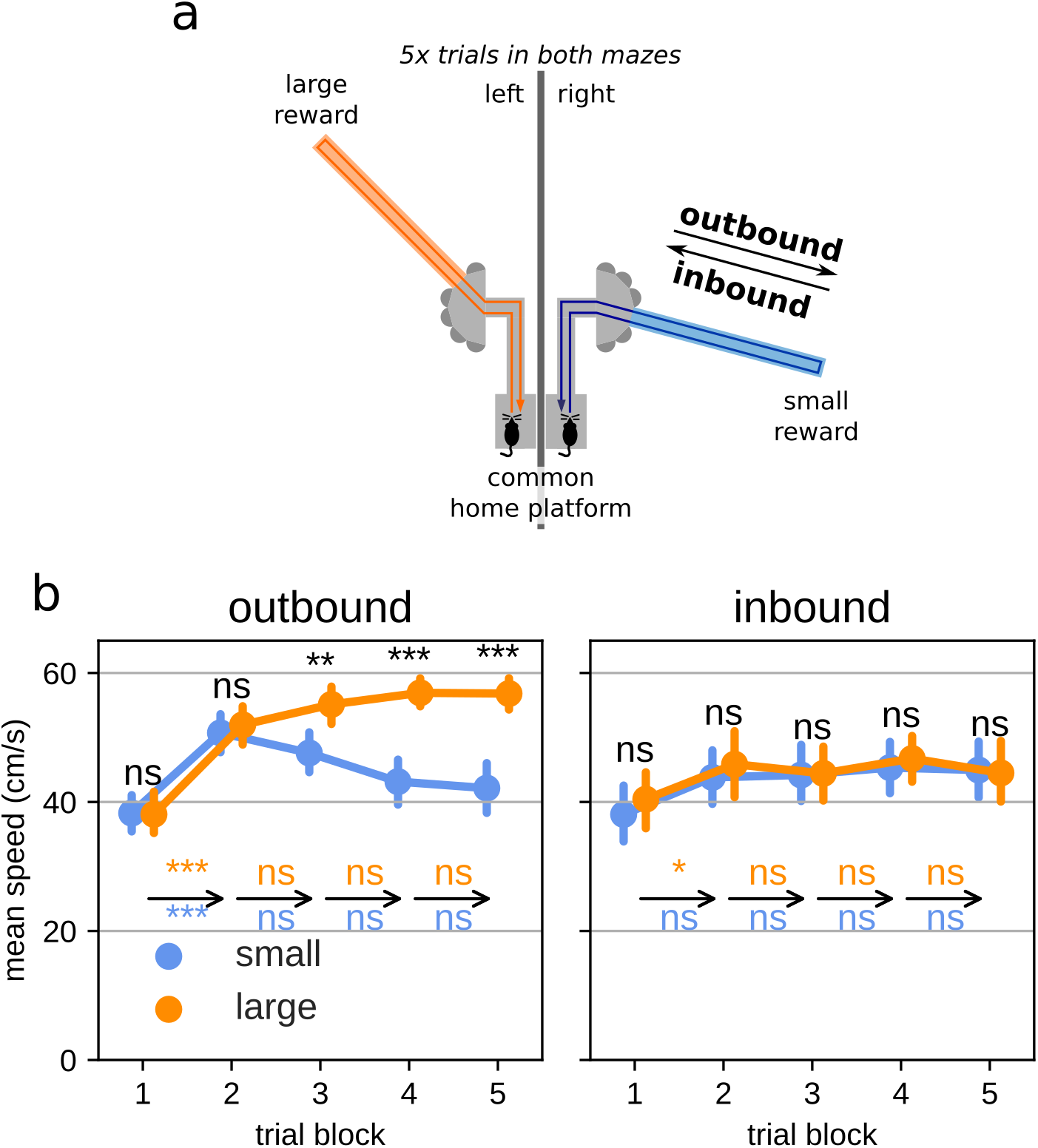
Fast learning of the reward-place associations during training. (a) Schematic representation of the apparatus. During training, rats learn to associate a small reward (blue) or a large reward (orange) with a specific arm in the left and right environment by alternating visits for five trials in each environments. Every day, the target arm location, the reward-environment associations and visiting order were pseudo-randomly assigned. (b) Average run speed over trial blocks separated by reward conditions and for journeys towards (outbound, left panel) or from (inbound, right panel) the reward location. Error bars indicate 95% bootstrapped confidence interval. Text annotations: arrows represent the trial-to-trial transitions annotated with the results statistical test for large (orange) and small (blue) reward. All pairwise comparisons were performed with Wilcoxon signed-rank test followed with Holm-Sidak correction for multiple tests. ***: p<0.001; **: p<0.01; *: p<0.05, ns: not significant.

To characterize the dynamics of the hippocampal spatial representation throughout the learning phase, we recorded cell activity from hippocampal subregion CA1 (**figure 2**). In a total of 37 separate sessions (median 7.0 sessions/animal, range 2-8 sessions/animal), we recorded from 758 place cells (median 18.0 cells/session, range 6-49 cells/session). For each cell, place fields were defined separately for outbound (home to reward platform) and in-bound (reward platform to home) run trajectories, and separately for small and large reward trials. Place cells had a median of 2.0 place fields (percentage of clusters with 1 field: 45.8%, 2 fields: 28.2%, 3 or more fields: 26.0%). Since the home location in the maze overlapped between run trajectories to/from the reward locations, we excluded place fields in home from the analyses below.

**Figure 2:**
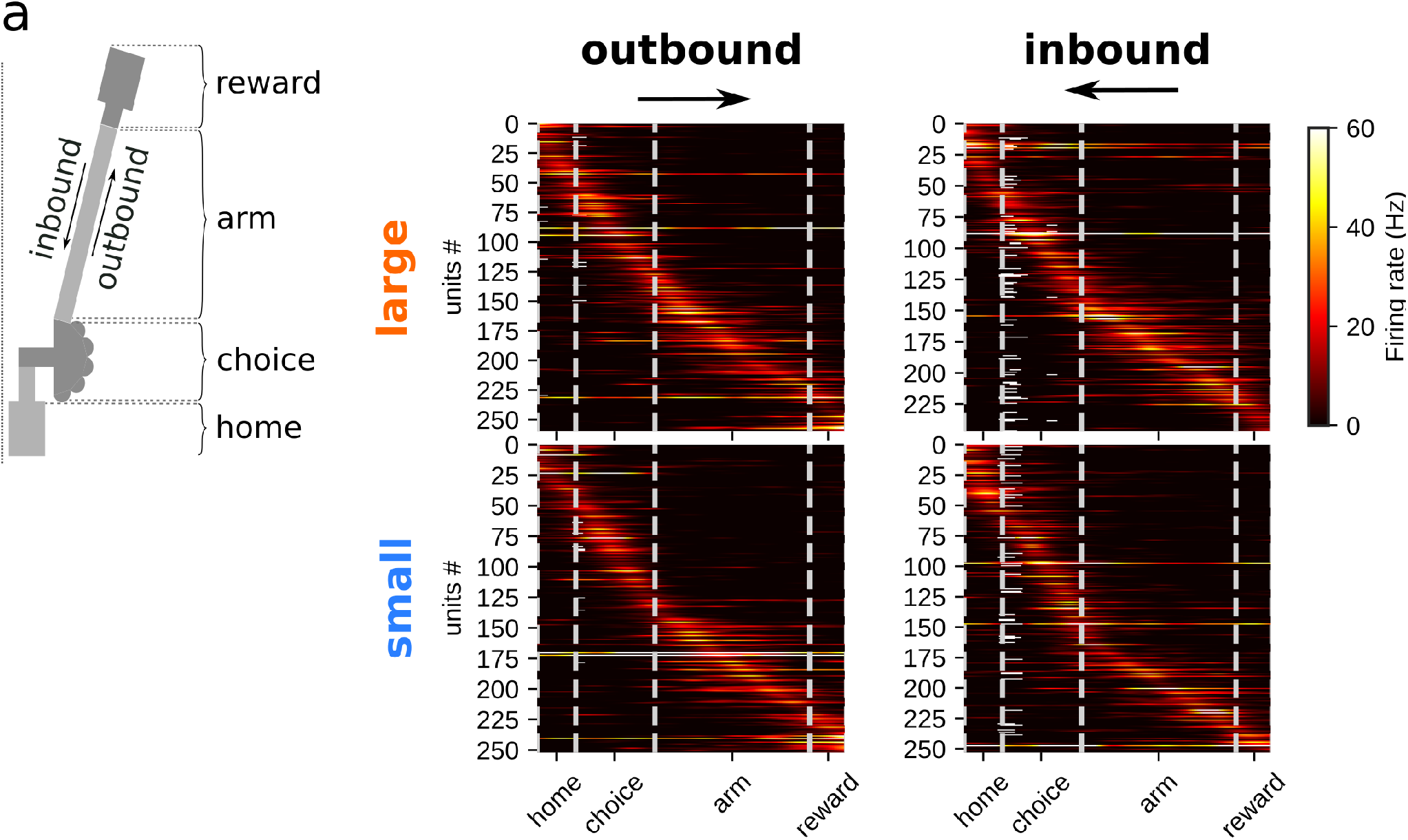
Spatial rate maps (a) Schematic representing the different maze segments in both environments. (b) Spatial rate maps for units active during run from all training sessions, ordered per peak rate locations and separated for runs on outbound (left column), inbound (right column) journeys, in the large reward environment (top row) and small reward environment (bottom row).

We first quantified the firing rate changes at the population level across trial blocks. Given the changes in run speed across trial blocks and the positive relation between speed and firing rate in hippocampal place cells (McNaughton, Barnes, and O’Keefe 1983), we corrected the in-field firing rates to remove the contribution of speed (**suppl. figure S1**, see Methods). Next, we constructed for each trial block a vector of in-field (speed-corrected) firing rates from all place fields and computed the spearman correlation coefficient for each pair of trial blocks (**figure 3a**). We observed that the population activity for trial blocks 2-5 are highly correlated (*r* in range 0.45-0.72), whereas the activity in the first trial block has systematically lower correlation with the other trial blocks (*r* in range 0.14-0.35). We then focused on the population vector correlations for consecutive trial block pairs (**figure 3b**). Across trial blocks, the correlation is lowest between trial blocks 1 and 2 (correlation coefficient *r* and **95%** bootstrapped confidence interval; *r*_1→2_=0.35 [0.24,0.46]) and highest between trials blocks 4 and 5 (*r*_4→5_=0.72 [0.68,0.76]). The population activity pattern is already highly similar between trial blocks 2 and 3 (*r*_2→3_=0.62 [0.56,0.67]). These results indicate that the hippocampal spatial representation quickly reorganizes after animals visited the target arm and experienced the associated reward size for the first time.

**Figure 3:**
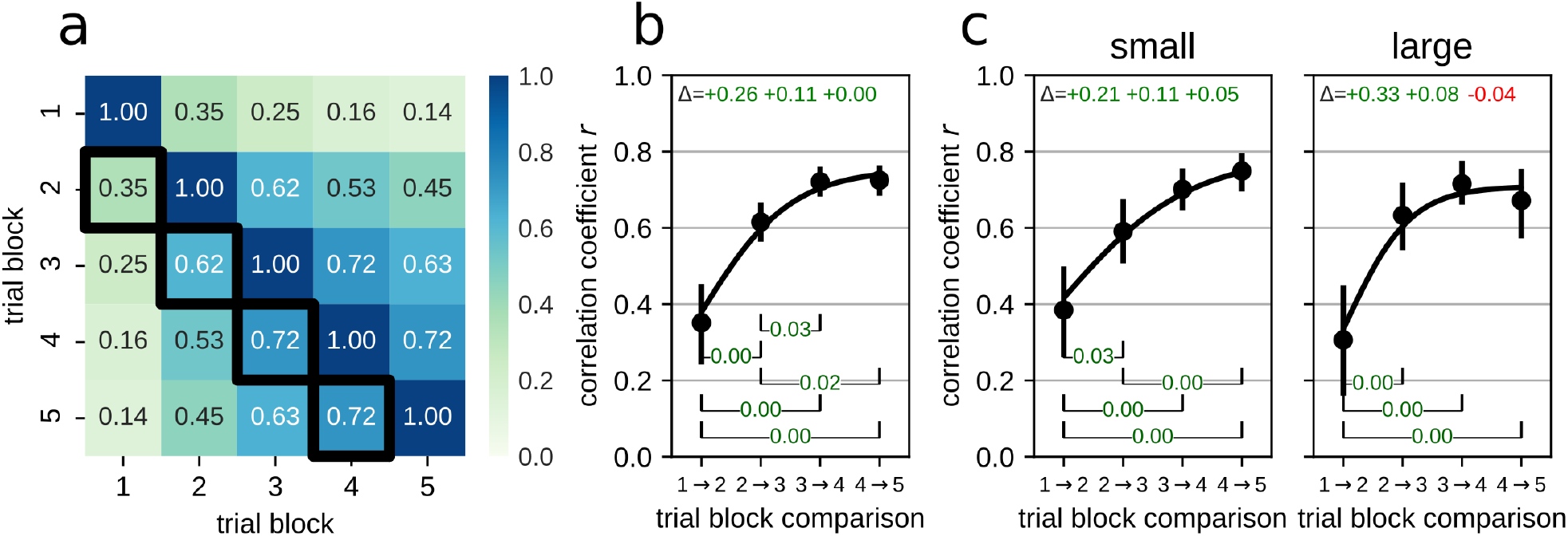
Trial-to-trial correlations show fast stabilization of the spatial code. (a) Pairwise correlations of the in-field firing rate vectors for all place fields and for all trial-block combinations. Note that pairwise correlations between trial blocks 2-5 are higher than correlations involving trial block 1. (b) Correlations of the in-field firing rate vectors for sequential trial block pairs (highlighted by black outlines in (a)). Error bars represent **95%** bootstrapped confidence interval. Black line represents fit of growth curve. Note that the correlation between trial blocks 1 and 2 is low, and the correlation in subsequent trial block pairs is high. Annotations at the top of the plot indicate consecutive pairwise changes in correlation. A one-way repeated measures analysis of variance by ranks (Friedman test) showed significant differences between trial-block pairs (statistic=37.44, p=3.7×10^−8^). Annotations at the bottom of the plot indicate p-value significant pairwise differences (Conover posthoc test with Holm-Sidak p-value correction). Coefficients and 95% confidence interval of the fitted growth curve: *β*_1_=0.75 [0.72,0.80], *β*_2_=0.99 [0.60,1.55], *λ*=1.32 [0.88,1.79]. (c) Correlations of the in-field firing rate vectors separately for small (left) and large (right) reward conditions. Annotations at the top of the plots indicate consecutive pairwise changes in correlation. For both reward conditions, Friedman test indicated significant differences between trial-block pairs (small: statistic=31.49, p=6.7×10^−7^; large: statistic=18.15, p=0.00041). Annotations at the bottom of the plots indicate p-value for significant pairwise differences (Conover posthoc test with Holm-Sidak p-value correction). Coefficients and 95% confidence interval of the fitted growth curve, small: *β*_1_=0.79 [0.74,0.86], *β*_2_=0.90 [0.58,1.35], *λ*=0.91 [0.56,1.36]; large: *β*_1_=0.71 [0.64,0.77], *β*_2_=1.12 [0.52,2.08], *λ*=1.86 [1.13,2.53].

We next looked at the trial-to-trial population-level activity changes separately for small and large reward instruction trials (**figure 3c**). The correlation between the first two trials is low for both small and large reward conditions (correlation coefficient and 95% bootstrapped confidence interval; small: *r*_1→2_=0.38 [0.27,0.50], large: *r*_1→2_=0.31 [0.16,0.50]). However, we observed a difference in the dynamics of the correlations across trials for the two reward conditions. Whereas for small reward the trial-to-trial correlation gradually increased towards the end of the instruction phase, for large reward we found a step-change between *r*_1→2_ and *r*_2→3_ and little further increase in later trials. We fitted the data with a sigmoid growth curve and compared the growth rate *λ* between reward conditions. As expected, growth rate *λ*_*large*_ was significantly larger than *λ*_*small*_ at *α* = 0.05 level (*λ*_*small*_ − *λ*_*large*_ 95% bootstrapped confidence interval: [−1.72,−0.12]).

These data suggest that in the large reward condition, place field activity abruptly reorganizes after the first trial and subsequently remains stable for the remainder of the trials. For the small reward condition, however, the reorganization and stabilization happens gradually over the 5 trials.

### Reorganization is accompanied by predominantly positive rate changes

To identify how the activity in individual place fields evolves over the instruction trials, we looked at the in-field rate changes between sequential pairs of trial blocks. The distribution of absolute rate changes between trial block 1 and 2 (|Δ*rate*_1→2_|) is significantly shifted to higher values as compared to distributions for other trial block pairs in particular for the large reward condition (**figure 4a**; two-sample Kolmogorov-Smirnov test for equal distributions |Δ*rate*_1→2_| vs |Δ*rate*_2→3_|; small: D=0.11 p*=0.0037, large: D=0.15 p*=6.3×10^−6^).

**Figure 4:**
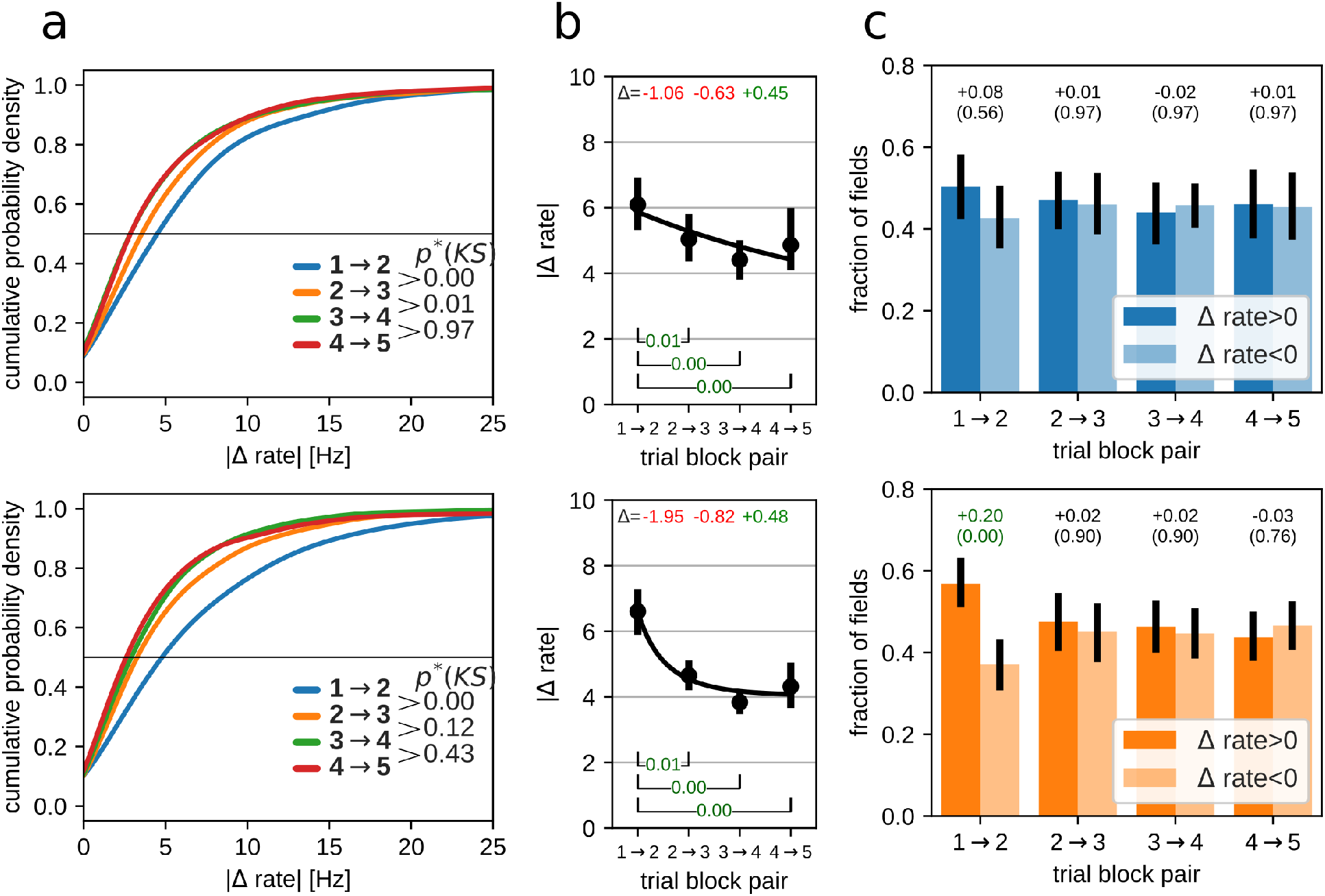
Trial-to-trial changes of in-field firing rate. Top row: small reward, bottom row: large reward. (a) Distributions of the magnitude of in-field rate changes |Δ*rate*| between pairs of trial blocks for all place fields in all sessions. p*(KS) is the Holm-Sidak corrected p-value for Kolmogorov-Smirnov test for equal distributions. (b) Average per-session magnitude of in-field rate changes for pairs of trial blocks. Error bars indicate 95% boot-strapped confidence interval. Text annotations at the top indicate the difference between consecutive trial block pairs. For both reward conditions, Friedman test indicated significant differences between trial-block pairs (small: statistic=20.51, p=0.00013; large: statistic=23.63, p=3×10^−5^). Annotations at the bottom of the plots indicate p-value for significant pairwise differences (Conover posthoc test with Holm-Sidak p-value correction). Coefficients and 95% confidence interval of the fitted growth curve, small: *β*_1_=0.25 [0.18,4.55], *β*_2_=−0.96 [−0.97,−0.29], *λ*=0.00 [0.00,1.18]; large: *β*_1_=4.04 [3.13,4.52], *β*_2_=−0.39 [−0.53,−0.29], *λ*=1.26 [0.44,3.27]. (c) Average fraction of fields with increasing (**dark**) or decreasing (**light**) in-field firing rate. Error bars indicate 95% bootstrapped confidence interval. Text annotations: top row, fraction difference between fields with increasing and decreasing in-field firing rate; bottom row: Holm-Sidak corrected p-value for Wilcoxon signed-rank test between increasing and decreasing rate fractions.

Consistent with the population vector analysis, the largest per-session absolute rate changes are observed from trial block 1 to 2 for small and large reward conditions (**figure 4b**). For subsequent trial block pairs, the absolute rate changes progressively declined, but at different rates for small and large reward.

We fitted the data with a sigmoid growth curve and compared the growth rate *λ* between reward conditions. The growth rate *λ*_*large*_ was considerably larger than *λ*_*small*_, although not significant at *α* = 0.05 level (*λ*_*small*_ − *λ*_*large*_ 95% bootstrapped confidence interval: [−3.22,0.29]).

We next looked at whether the absolute rate changes reflect both increases and decreases of in-field firing rate. For this, we computed the per-session fractions of place fields with rate increase or decrease (**figure 4c**). For large reward but not small reward, we found a significant bias towards rate increases from trial block 1 → 2 (Wilcoxon signed-rank test between increasing and decreasing rate fractions, small: W=235.50, p*=0.58; large: W=69.50, p*=0.0011). For other trial-block pairs, we observed an equal proportion of rate increase and rate decrease.

### A subset of place fields emerge or vanish after initial trials

We noted that a subset of place fields were not active (i.e. zero in-field rate) during one or more trials. We wondered if zero-rate trials more likely occur in the first trial(s) (i.e. emerging fields) and last trial(s) (i.e. vanishing fields), rather than being randomly distributed across trials as in the case of a noisy signal.

We tested whether the fraction of fields with *n* zero-rate trials on the trials *[t…t+n-1]* is higher than random chance (computed as the probability of drawing *n* trials out of 5). For example, for all fields with two zero-rate trials, we counted the fraction of fields for which the first two trials were zero-rate trials and divided this by the expected fraction under the assumption of random distribution of the zero rate trials across all the 5 instruction trials.

We found a clear overrepresentation of place fields that were not active in the first 1-3 trials, but became active afterwards (emerging fields), and place fields that were initially active but then disappeared after trials 1-3 (vanishing fields) (**figure 5a**).

**Figure 5:**
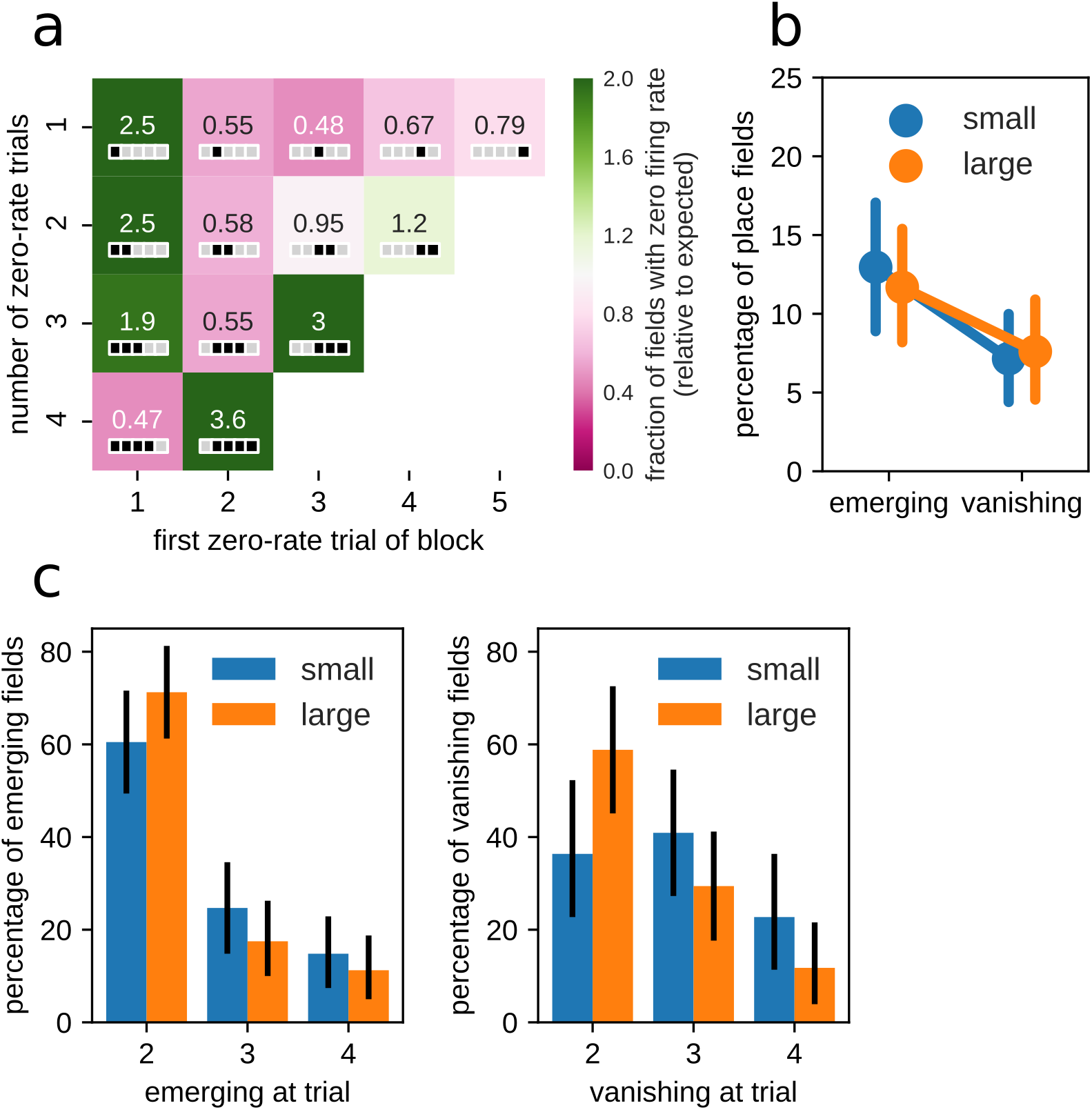
A subset of place fields emerge or vanish after the initial three trial blocks. (a) The number of place fields with a string of *n* zero-rate trials starting at trial block *t* was expressed relative to the expected number place fields with *n* zero-rate trial blocks (assuming a uniform distribution of zero-rate occurrences across the 5 trial blocks). For example, the value of 2.5 at (1,1) indicates that it is more likely than expected that a single zero-rate trial occurs in the first trial block. For each cell in the matrix, the inset depicts the pattern of zero-rate trial blocks (black/gray squares represent zero-rate/non-zero-rate trial blocks respectively). (b) Percentage of emerging and vanishing place fields for small and large reward conditions. Note that there are significantly more emerging place fields than vanishing place fields for both reward conditions, but there is no difference between small and large reward. (c) Distribution of trial blocks at which place fields emerge (left) or vanish (right). For both small and large reward conditions, place fields emerge predominantly in trial block 2. For the large reward condition, place fields predominantly disappear in trial block 2. For the small reward condition, the distribution is not statistically different from uniform, and the highest percentage of place fields disappear in trial block 3.

For both small and large reward condition there was a signficantly larger fraction of emerging place fields than vanishing place fields (**figure 5b**; Wilcoxon test for equal fractions of emerging and vanishing fields, small: statistic=77.00, p=0.012; large: statistic=98.00, p=0.029). We next looked at when place fields emerged or vanished. Emerging fields did not appear uniformly on trial blocks 2-4 (Chi-square test for null hypothesis of uniform distribution, small: statistic=28.07, p=8×10^−7^; large: statistic=52.22, p=4.6×10^−12^), but rather most emerging fields appeared after the first trial block (**figure 5c**). This pattern of field emergence was the same for small and large reward. For vanishing fields in the large reward condition, a similar bias towards disappearing after the first trial was observed (**figure 5c**, Chi-square test, statistic=17.29, p=0.00018). For small reward, however, the distribution is not statistically different from uniform (Chi-square test, statistic=2.36, p=0.31), and the highest percentage of vanishing place fields disappear in trial block 3.

Overall, these results indicate that the appearance of place fields occurs rapidly after the first trial. Furthermore, the slower dynamics of place field disappearance in the small reward condition could contribute to the overall slower stabilization of the small reward spatial representation.

The question arises if the emerging and vanishing fields explain the dynamics of the spatial representation changes over trial blocks. We repeated the population vector correlation and rate change analysis, but now excluding emerging and vanishing fields (figures S2 and S3). Overall, the the trial-to-trial dynamics of the population vector correlations and rate changes persist and are thus not solely explained by the emergence or disappearance of place fields.

### Place cell activity in SWRs is modulated by experience and reward amount

Periods of consumatory behavior during training are characterized by the reactivation of hippocampal place cell assemblies (replay) (Buzsáki 2015). Hippocampal replay activity is modulated by both experience and reward (O’Neill et al. 2008; Ambrose, Pfeiffer, and Foster 2016). Previously, we showed that in the dual reward-place association task hippocampal replay during the instruction phase is strongly biased towards the large reward environment (Michon et al. 2019). In the following analyses, we ask how replay activity is modulated by experience across the trial blocks during learning of the reward-place associations and how replay is related to the changes in the spatial representations shown above.

First, we examined the occurence of sharp-wave ripples (SWRs), electrophysiological markers of putative replay in the local field of CA1. The large majority of SWRs occured while the animal was consuming the large reward and only very few occured when consuming the small reward or at other locations (**figure 6a**, mean±sem (percentage±sem) number of SWRs per session: large reward platform, 78.6±5.1 (88.0%±1.4); small reward platform, 2.1±0.7 (2.1%±0.6)).

**Figure 6:**
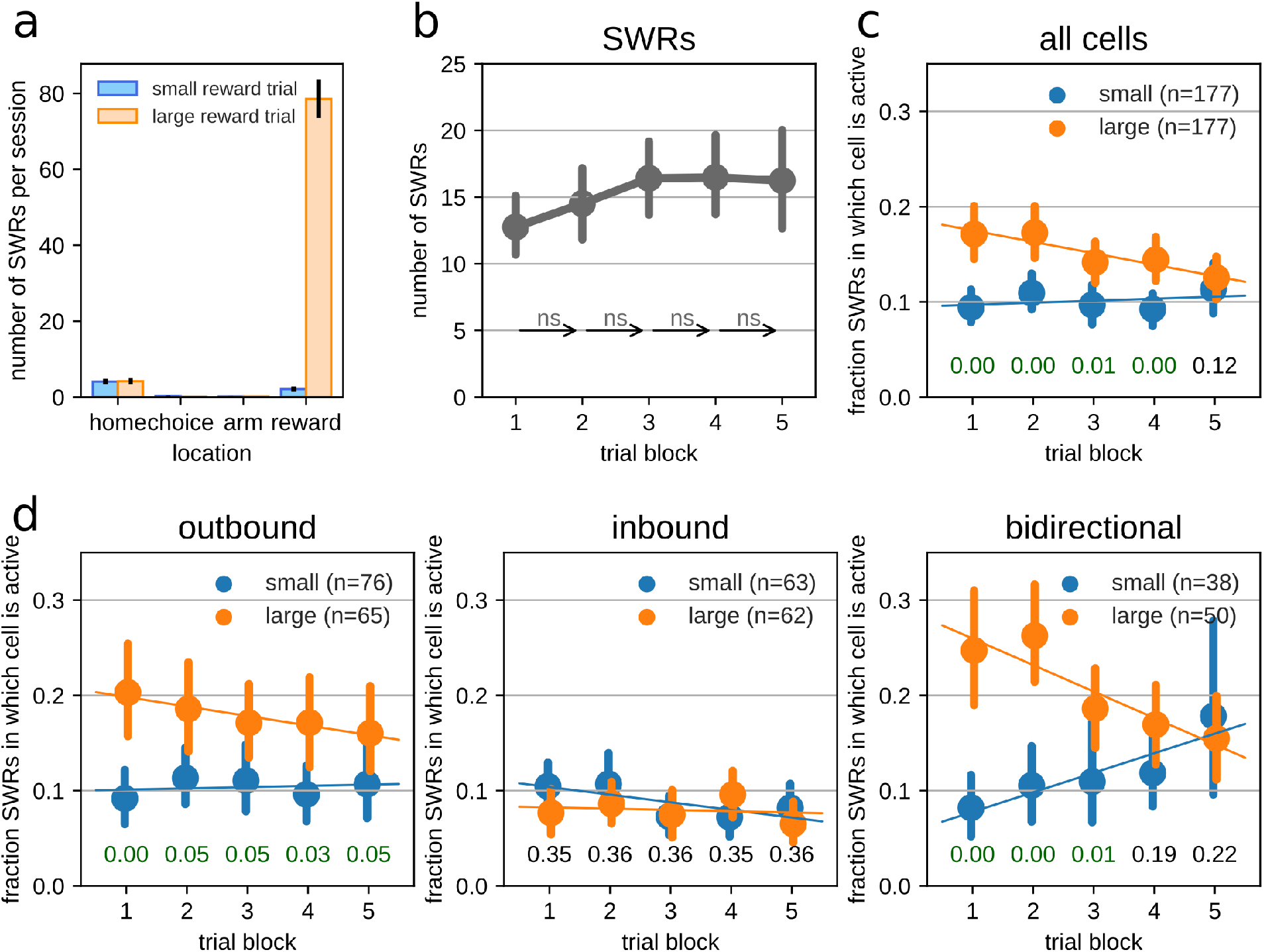
Place cell activity during SWRs changes over trial blocks. (a) Average number of SWRs per session as a function of location in the large reward and small reward environments. Note that the large majority of SWRs occur when rats visit the large reward platform. (b) The number of SWRs per trial block. Arrows represent the trial-to-trial transition annotated with the significance level. ns: not significant. (c) The fraction of SWRs during which place cells are active, separated for cells with fields in the small and large reward environments. Text annotations at the bottom indicate the corrected p-value for Mann-Whitney U tests between large and small reward conditions. Error bars indicate 95% bootstrapped confidence intervals. Lines represent least squares linear fit. (d) Same as (c) for place cells with fields in the outbound direction (left), in the inbound direction (middle) or in both directions (right).

Across trials, the number of SWRs trended higher for later trial blocks (**figure 6b**; least-square linear regression, slope=0.89 events per trial block, R=0.15, p=0.065) without increase in total time spent immobile (**supp figure S4**). Pairwise comparison of consecutive trial blocks did not reveal significant trial-to-trial changes in the number of SWRs (Wilcoxon signed-rank test, trial block 1 → 2: statistic=160.00, p*=0.51; trial block 2 → 3: statistic=134.50, p*=0.26; trial block 3 → 4: statistic=217.50, p*=1; trial block 4 → 5: statistic=216.00, p*=1).

We next asked how individual place cells change their activity within SWRs over trial blocks. For this we computed the proportion (*p*) of SWRs in which a place cell fired at least one spike. We only considered contextual place cells with place fields on the path to either large or small reward (*p*_*small*_ and *p*_*large*_) and excluded cells with fields on both paths or with fields in the common home. Overall, cells with fields on the path to the large reward were active in a higher proportion of SWRs then cells with fields on path to small reward (**figure 6c**). However, *p*_*large*_ decreased across trials (least-square linear regression, slope=−0.01 per trial block, R=−0.10, p=0.0028), such that *p*_*large*_ and *p*_*small*_ significantly differed in all trial blocks except the last (Mann-Whitney U test for each trial block with Holm-Sidak multiple test correction of p-values, trial block 1: U=12019.50, p*=0.00025, trial block 2: U=12861.50, p*=0.0046, trial block 3: U=13123.00, p*=0.0071, trial block 4: U=12722.50, p*=0.0037, trial block 5: U=14563.50, p*=0.12).

On linear tracks, place cell activity is largely directional (McNaughton, Barnes, and O’Keefe 1983) and in hippocampal replay events the reactivated place cells often represent either the outbound or inbound spatial map, giving rise to forward and reverse replay sequences (Foster and Wilson 2006; Diba and Buzsáki 2007; Davidson, Kloosterman, and Wilson 2009). We next looked at the activity in SWRs separately for unidirectional and bidirectional place cells. For directional place cells, *p*_*large*_ is significantly higher than *p*_*small*_ across all trial blocks only for cells that are active in the outbound run (i.e. towards the reward), but not for cells active only in the inbound run (**figure 6d**, Mann-Whitney U test for each trial block with Holm-Sidak multiple test correction of p-values, outbound place cells, trial block 1: U=1573.00, p*=0.00035, trial block 2: U=2007.00, p*=0.046, trial block 3: U=1958.00, p*=0.046, trial block 4: U=1906.00, p*=0.033, trial block 5: U=1978.50, p*=0.046; inbound place cells, trial block 1: U=1705.00, p*=0.35, trial block 2: U=1765.00, p*=0.36, trial block 3: U=1877.50, p*=0.36, trial block 4: U=1679.00, p*=0.35, trial block 5: U=1738.50, p*=0.36;). There was a negative trend in *p*_*large*_ across trial blocks, but this was not significant (least-square linear regression, slope=−0.01 per trial block, R=−0.07, p=0.18). For place cells active in the inbound direction in the small reward environment, *p*_*small*_ modestly but significantly decreased across trial blocks (least-square linear regression, slope=−0.01 per trial block, R=−0.11, p=0.049). Interestingly, the largest changes across trial blocks was seen for bidirectional cells with place fields in both outbound and inbound directions (**figure 6d**). We found an increase of *p*_*small*_ and decrease of *p*_*large*_ across trial blocks (least-square linear regression, *p*_*large*_: slope=−0.03 per trial block, R=−0.22, p=0.00053; *p*_*small*_: slope=0.02 per trial block, R=0.16, p=0.026), with a significant difference between *p*_*small*_ and *p*_*large*_ in the first three but not the last two trial blocks (Mann-Whitney U test for each trial block with Holm-Sidak multiple test correction of p-values, bidrectional place cells, trial block 1: U=489.00, p*=0.00021, trial block 2: U=487.00, p*=0.00021, trial block 3: U=605.50, p*=0.0053, trial block 4: U=797.50, p*=0.19, trial block 5: U=857.50, p*=0.22).

### Replay activity is modulated by experience and reward amount

To quantify replay events during periods of immobility, we used a bayesian neural decoding approach to detect spatial representations during SWR bursts. An encoding model was constructed from the unsorted spatially-tuned neuronal activity recorded in the hippocampus during the runs to and from the reward site. Cross-validation analysis was used to assess the precision with which the model could estimate the animal’s position. Sessions with a 75 percentile decoding error above 30 cm were excluded, leaving 29 sessions from 6 animals.

Previously, we used an encoding model that incorporates hippocampal activity from all trials (trial-average model) (Michon et al. 2019). That analysis showed a strong bias for reverse replay of the outbound trajectory towards the large reward. Here, we investigated how the expression of replay changes across trial blocks.

The trial-average model assumes that the spatial tuning of hippocampal place cells is constant and in that case reduces error in the model due to variations in spiking. Since we showed that place field activity systematically changes across trials block with different dynamics for small and large reward conditions, the trial-average model may not be the best model to analyze replay events on individual trial blocks. In particular for the first trial block in which place field activity has the lowest correlation with other trial blocks. For this reason, we performed analysis of replay with per-trial block encoding models (single-trial model) (**figure 7** and **suppl figure S5**). The results using the trial-average encoding model are described in the supplemental data (**suppl figure S6**).

**Figure 7:**
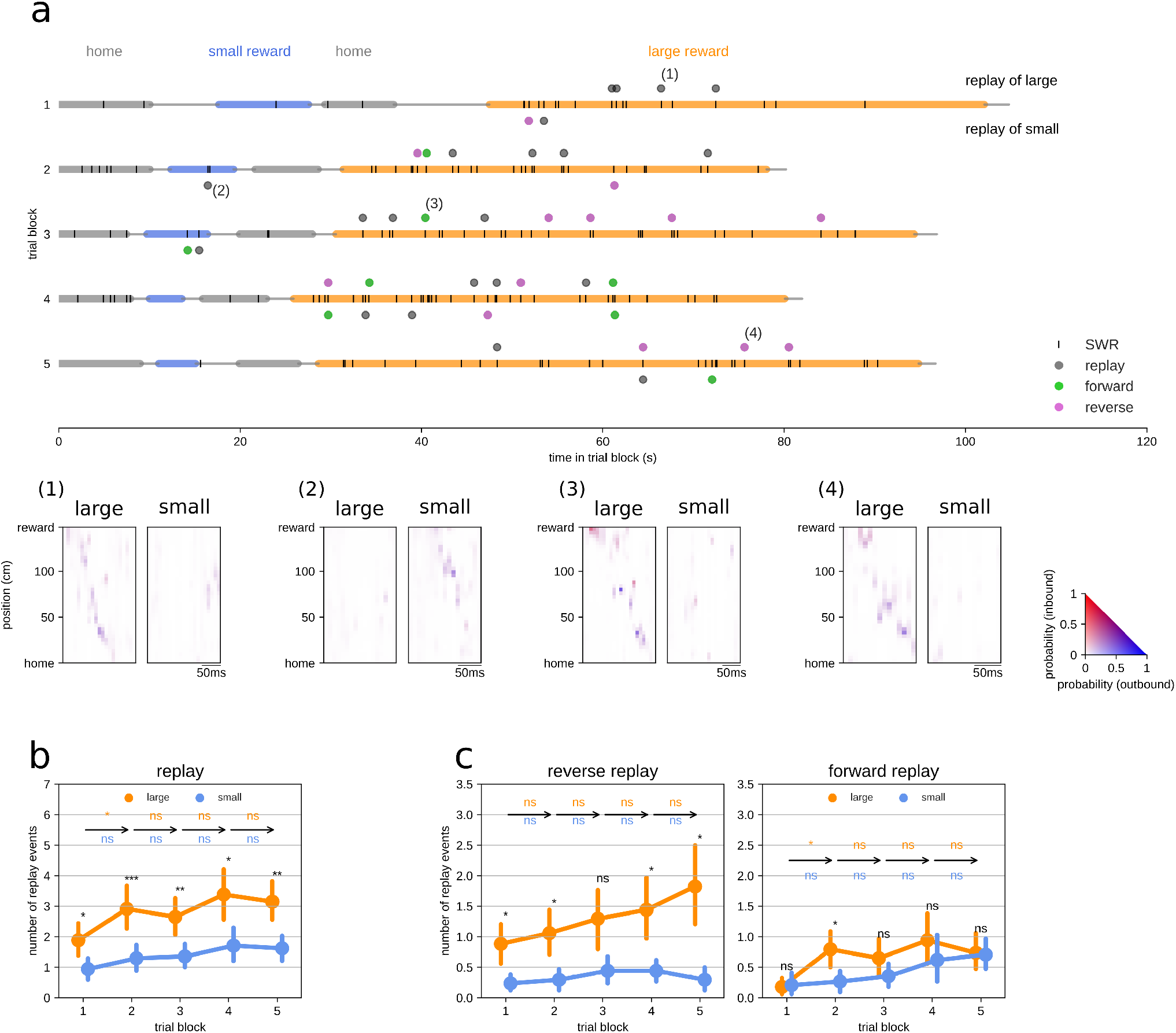
Changes in replay over training and between reward conditions. (a) Examples of identified replay events in one session. (b) Number of 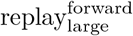 and 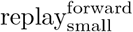 events across trial blocks. (c) Number of reverse (left) and forward (right) replay events across trial blocks. Error bars indicate **95%** bootstrapped confidence interval. Text annotations: arrows represent the trial-to-trial transitions annotated with the significance level separately for the two reward groups. For each trial block, significance level for the difference between the two reward groups is indicated above the data points. All pairwise comparisons were performed with Wilcoxon signed-rank test followed by Holm-Sidak correction for multiple tests. ***: p<0.001; **: p<0.01; *: p<0.05, ns: not significant.

Trajectory replay was characterized using a weighted isotonic regression fit on the decoded positions during SWR bursts (Michon et al. 2019). Only events with a p-value < 0.05 were considered significant trajectory replays. Note that since the large majority of SWRs and replay events occured at the large reward platform, by definition most replay_small_ are remote and replay_large_ are predominantly local (Davidson, Kloosterman, and Wilson 2009; Karlsson and Frank 2009).

Next, we analyzed how the number of replay events varied across trial blocks. Overall, the number of replay_large_ events is higher than the number of replay_small_ for every trial block (Wilcoxon signed-rank test: trial block 1: W=94.00, p*=0.014; trial block 2: W=16.50, p*=6.5×10^−5^; trial block 3: W=68.00, p*=0.0034; trial block 4: W=97.00, p*=0.014; trial block 5: W=60.00, p*=0.0025). We find a gradual and significant increase in the number of replay events that express trajectories in the small and large reward environments (least-square linear regression, replay_large_: slope=0.30 events per trial block, R=0.20, p=0.0076; replay_small_: slope=0.18 events per trial block, R=0.19, p=0.013). In addition, we observed a small significant jump in replay_large_ events from trial block 1 → 2 (Wilcoxon signed-rank test: W=128.00, p*=0.014) (**figure 7b**).

Hippocampal reactivation of place cells in the opposite order as that observed during behavior (reverse replay) has been previously shown to be uniquely modulated by reward size in contrast to reactivation in the same sequential order as during behavior (forward replay) (Ambrose, Pfeiffer, and Foster 2016). We thus further explored the relationship between reverse and forward replay activity (characterized as having a decoded direction opposite or similar to that of the trajectory fit respectively), experience and reward size (**figure 7c**).

We found that the number of 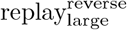 events but not 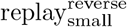 events significantly increased across trial blocks (least-square linear regression, 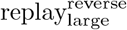: slope=0.226 events per trial block, R=0.21, p=0.0053; 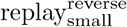: slope=0.03 events per trial block, R=0.07, p=0.37) without sudden trial block-to-trial block changes. The average occurence of 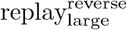 events was consistently higher than 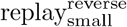 across all trial blocks (Wilcoxon ranked-test, trial block 1: W=90.50, p*=0.0092; trial block 2: W=80.00, p*=0.0082; trial block 3: W=88.50, p*=0.0092; trial block 4: W=42.00, p*=0.00052; trial block 5: W=36.50, p*=0.0004).

The number of forward replay events increased for both reward conditions throughout the trial blocks (least-square linear regression, 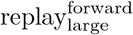: slope=0.13 events per trial block, R=0.19, p=0.014; 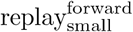: slope=0.14 events per trial block, R=0.25, p=0.0011). The average occurence of 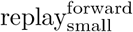 did not show large changes on a trial block-to-trial block basis. In contrast, the average number of 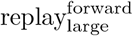 events strongly increased from trial block 1 → 2 (Wilcoxon signed-rank test, trial block 1 → 2: W=130.50, p*=0.015). As a result, the difference between the number of 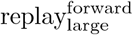 and 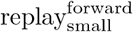 events was significant on trial block 2 (Wilcoxon signed-rank test: W=89.50 p*=0.024).

Note, interestingly the dynamics of replay occurence reported above were similar to those observed with the average-encoding model for all but the 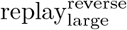 events (**suppl figure S6**). Further, the number of detected 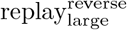 events was consistantly higher when using single-trial encoding models close to that during which the events were decoded (**suppl figure S5**), suggesting that reverse replays may be more tightly related to single experiences than forward replays.

Taken together, these results suggest that subsets of hippocampal replay are differently modulated by reward and experience. Reverse replay activity was strongly modulated by reward size and forward replay activity increased with experience with a distinct reward effect in the second trial block following the large changes in the place field activity that we showed above.

## Discussion

Our aim was to assess the influence of reward value during learning on hippocampal representations in a paradigm where rats repeatedly learn different reward-place associations in a familiar setting. Thanks to the structure of the paradigm, we could assess the changes in hippocampal CA1 neuronal representations, both during active exploration and during consummatory state, that are associated with learning and for experiences associated with large or small reward amount in parallel.

Overall, active exploration of the two environments was accompanied by the fast reorganization of hippocampal place cells firing over the course of a few training trials. The changes in hippocampal spatial representations were the results of both the appearance and disappearance of place fields and the modulation of firing rate. These observations are consistent with the body of literature showing that changes in an environment (Kentros et al. 1998; Anderson and Jeffery 2003), exposure to a different environment (Lever et al. 2002; Leutgeb et al. 2004, 2005) and spatial learning (Frank, Stanley, and Brown 2004; Karlsson and Frank 2008; Dupret et al. 2010) induces a fast partial remapping of the spatial representation in the hippocampus (Anderson and Jeffery 2003), with the modulation of within-field firing rate (rate remapping) and the formation of new fields and disappearance of existing ones (global remapping) for different subgroups of neurons (Leutgeb et al. 2004, 2005; Latuske et al. 2018). Consistent with the accumulation of field activity on novel or changed goal locations (Fyhn et al. 2002; Dupret et al. 2010; Danielseon et al. 2016) the ratio emerging fields / vanishing fields was highest at the home and reward locations. However, similarly to what has been previously reported (Duvelle et al. 2019) we did not observe a difference in the final representations of the environments associated with large and small rewards. This suggests that salient goal locations are overrepresented in dorsal CA1 place code during learning, regardless of their associated reward value.

Conformally with the idea that reward value enhances learning speed (Wolosin, Zeithamova, and Preston 2012; Igloi et al. 2015; Miendlarzewska, Bavelier, and Schwartz 2016), the main difference between reward conditions was a faster reorganization and stabilization of the spatial map for the large reward environment. While most of the changes occurred between the first two trials in the large reward environment, with an overall increase in within-field firing rate associated to a higher turn over of active fields, the spatial representation of the small reward environment changed more progressively throughout learning.

The rapid reorganization of hippocampal representations, and in particular the formation of novel place fields, has been suggested to be at least in part due to changes in synaptic plasticity (Sheffield, Adoff, and Dombeck 2017). Associated reward amount may have accelerated the changes in spatial representation in CA1 via the, not mutually exclusive, modulation of dopamine and noradrenaline release. On the one hand, dopamine release increases with reward value and novelty (Tobler, Fiorillo, and Schultz 2005; Shohamy and Adcock 2010), and its release to the dorsal hippocampus enhances spatial learning (Kempadoo et al. 2016; Retailleau and Morris 2018). Dopamine enhances excitability and facilitates synaptic plasticity (Hansen and Manahan-Vaughan 2014; Rosen, Cheung, and Siegelbaum 2015; Sheffield, Adoff, and Dombeck 2017) after exposure to a salient experience [Retailleau2018]. Moreover, novelty and learning-induced changes in hippocampal spatial representation are abolished in the absence of dopamine activation (Tran et al. 2008; Retailleau and Morris 2018). On the other hand, salience experiences also upregulate noradrenaline activity (Izquierdo and Medina 1997; Roozendaal and McGaugh 2011). Further, adrenoreceptors activation in the hippocampus increases CA1 neurons excitability (Bacon, Pickering, and Mellor 2020), promotes synaptic plasticity and memory encoding (Lemon et al. 2009; Sara 2009), and is required for novelty-induced changes in hippocampal spatial code (Tanila 2001). Finally, inhibition of the locus coeruleus projections to the hippocampus, known to release both dopamine and noradrenaline (Kempadoo et al. 2016; Ranjbar-Slamloo and Fazlali 2020), impairs the accumulation of fields near newly rewarded locations (Kaufman, Geiller, and Losonczy 2020). The exact contribution of the neuromodulatory systems to the formation and the reorganization of hippocampal spatial maps require further investigation.

We also monitored replay activity during training using single-trial place activity during run as encoding model to control for the influence of the changes in representation throughout learning. The large majority of events occurred while the animals were consuming the large reward, possibly biasing our observations of replays towards events representing the current location of the animal (Davidson, Kloosterman, and Wilson 2009). Nevertheless, we did observe remote replays of the small reward environment (Karlsson and Frank 2009) and our observations are consistent with previous reports (Singer and Frank 2009; Ambrose, Pfeiffer, and Foster 2016). The dynamic of trajectory events occurrence during training resembled that of the changes in spatial representation during exploration, with a fast increase for replays of the large environment from the first trial block to the second one, and a more progressive increase throughout training for replays of the small reward environment. These results are consistent with the fact that replay activity is dependent on experience (O’Neill et al. 2008; Silva, Feng, and Foster 2015). However, the occurrence of subsets of events followed different dynamics. On the one hand, consistent with (Ambrose, Pfeiffer, and Foster 2016) we found that trajectory replay of the reverse order than observed during behavior was strongly increased by reward amount from the first trial onward; yet, it also progressively increased throughout training, suggesting a combined effect of both reward and experience on reverse replay activity. On the other hand, the dynamics of forward replay activity for both reward conditions followed that observed for the reorganization of the spatial map, indicating that it was modulated by experience, but with a minor influence of reward amount in the present paradigm (Ambrose, Pfeiffer, and Foster 2016). Moreover, the number of detected reverse replay of the large reward environment was found to be more sensitive to the encoding model used for decoding. We found a higher number of identified events when using the run activity that occured close to the time of the events occurence which suggest that reverse replay corresponds to the reactivation of single-experience related patterns of CA1 activity, more so than forward events.

Overall, replay activity became more strongly biased towards reactivation of directional trajectories with an overrepresentation of reverse replays of the large reward environment. Given that a majority of decoded spatial trajectories initiated at the reward locations and spaned the arm towards home (Michon et al. 2019), this suggests that with experience, the journeys leading to the large reward were more frequently replayed. This observation is further corroborated by the fact that the contribution of units active on both running trajectories in the small reward conditino decreased during learning while the contribution of individual units active on journeys towards the large reward was more elevated than those active in the small reward environment. Taken together, these results suggest that the reward-related enhancement of hippocampal replay activity is finely tuned towards behaviourally relevant experiences, in this case, the journeys leading to the large reward, and is learning-dependent.

Finally, the differences in the dynamics of reverse and forward replays occurrence point to the fact that, at least partially, different mechanisms contribute to the generation of such events and support the hypothesis that these subsets of hippocampal replay may play different roles for memory processes (Diba and Buzsáki 2007; Carr, Jadhav, and Frank 2011; Joo et al. 2018; Ólafsdóttir, Bush, and Barry 2018; Xu et al. 2019). The tighter bound of reverse replays, which can be observed after a unique lap (Foster and Wilson 2006), with single experiences and their unique modulation by reward (Ambrose, Pfeiffer, and Foster 2016) suggest that their generation may be more strongly driven by external inputs, such as from cortical areas (Ji and Wilson 2007; Rothschild, Eban, and Frank 2017) or reward responsive neurons present in the VTA (Gomperts, Kloosterman, and Wilson 2015). It also suggests that reverse replay enhances reward-related learning by retroactively strengthening cell assemblies active prior to obtaining a larger reward (Ambrose, Pfeiffer, and Foster 2016). Forward replays have been hypothesized to contribute to several memory processes such as memory consolidation and evaluation of future options (Pfeiffer and Foster 2013; Silva, Feng, and Foster 2015; Joo et al. 2018), both of which require prior encoding of a memory traces. Our observations that forward replays are more strongly related to the experience-dependent formation and modification of hippocampal spatial representations are consistent with this view. Further investigations will be needed to decipher the roles of subsets of hippocampal replay to memory, for example via their specific online manipulation during behavior (Deng et al. 2016; Ciliberti, Michon, and Kloosterman 2018).

## Acknowledgments

F.K. is funded through Flemish Research Project FWO G0D7516N and KU Leuven C1 grant C14/17/042.

## Author Contributions

Conceptualization, F.M. and F.K.; Formal Analysis, F.M., E.K. and F.K.; Investigation, F.M. and J.S.; Writing - Original Draft, F.M. and F.K; Writing - Review & Editing, F.M., J.S. and F.K; Visualization, F.M. and F.K.; Supervision, F.K.; Funding Acquisition, F.K.

## Declaration of Interests

The authors declare no competing interests.

## Supplemental Information

### Speed correction of in-field firing rates

**Figure S1:**
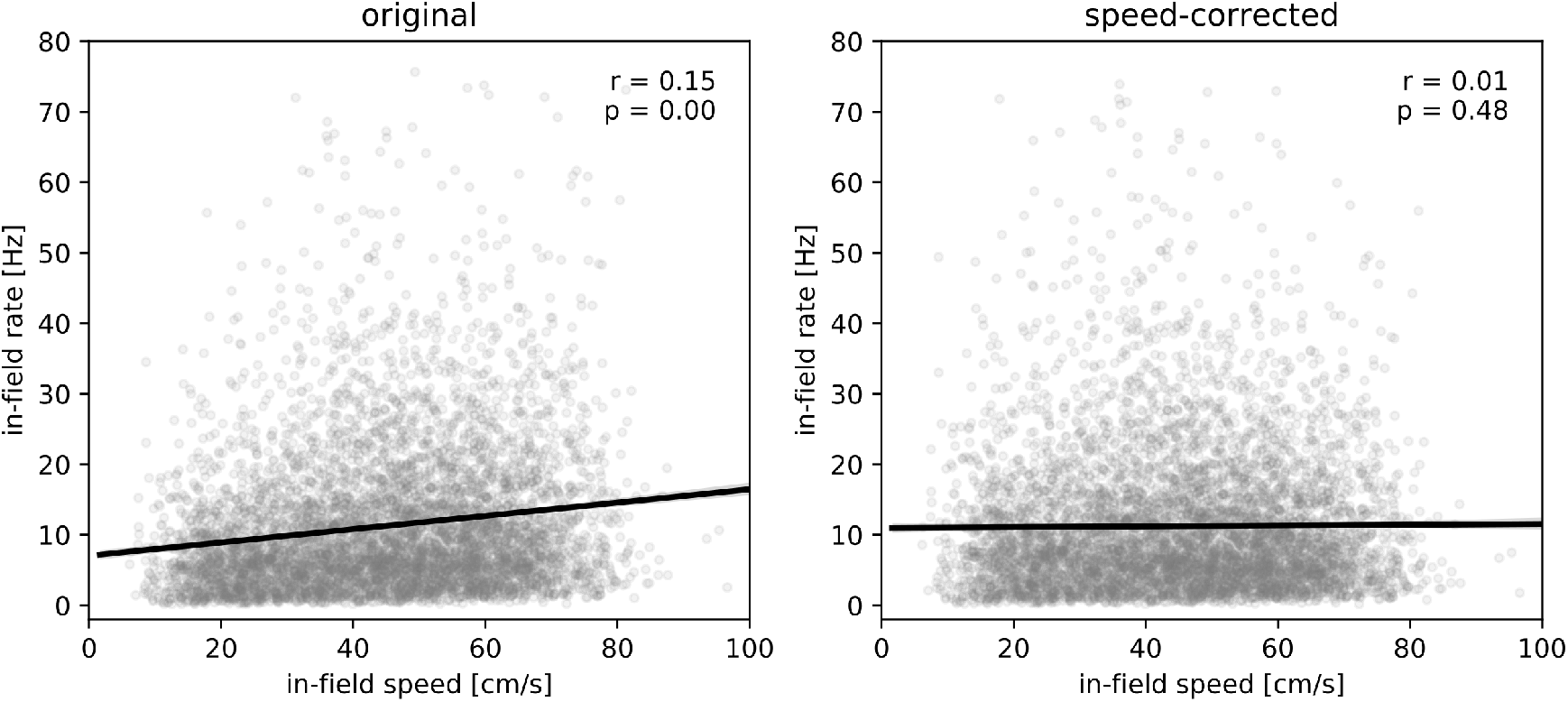
Speed correction of in-field firing rates. (a) Prior to correction, in-field firing rate is weakly but significantly correlated to running speed in the field. Black line indicates linear fit. Each dot represents the rate and run speed for a single place field on a single trial. Note that data samples for which the in-field firing rate is zero are not shown and are also not included in the correction procedure. (b) After correction, no significant correlation between in-field firing rate and run speed remains.

### Trial-to-trial correlations and rate changes without emerging/vanishing place fields

**Figure S2:**
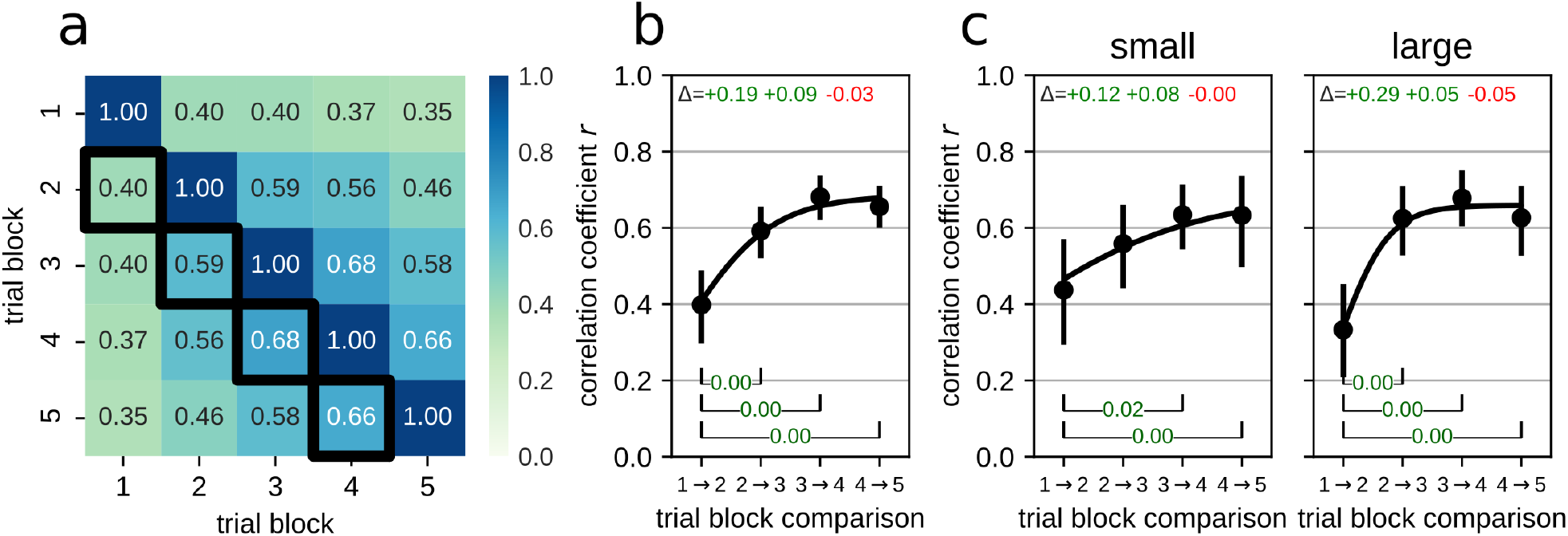
Trial-to-trial correlations show fast stabilization of the spatial code. Same analysis as in figure 3, but excluding emerging and vanishing fields. (a) Pairwise correlations of the in-field firing rate vectors for all place fields and for all trial-block combinations. Note that pairwise correlations for trial blocks 2-5 are higher than correlations involving trial block 1. (b) Correlations of the in-field firing rate vectors for sequential trial block pairs (highlighted by black outlines in (a)). Error bars represent **95%** bootstrapped confidence interval. Black line represents fit of growth curve. Note that the correlation between trial blocks 1 and 2 is low, and the correlation in subsequent trial block pairs is high. Annotations at the top of the plot indicate consecutive pairwise changes in correlation. A one-way repeated measures analysis of variance by ranks (Friedman test) showed significant differences between trial-block pairs (statistic=26.40, p=7.9×10^−6^). Annotations at the bottom of the plot indicate the p-value of significant pairwise differences (Conover posthoc test with Holm-Sidak p-value correction) indicated that *r*_1→2_. Coefficients and 95% confidence interval of the fitted growth curve: *β*_1_=0.69 [0.64,0.74], *β*_2_=0.68 [0.39,1.16], *λ*=1.38 [0.85,1.79]. (c) Correlations of the in-field firing rate vectors separately for small (left) and large (right) reward conditions. Annotations at the top of the plot indicate consecutive pairwise changes in correlation. For both reward conditions, Friedman test indicated significant differences between trial-block pairs (small: statistic=9.92, p=0.019; large: statistic=20.69, p=0.00012). Annotations at the bottom of the plot indicate the p-value of significant pairwise differences (Conover posthoc test with Holm-Sidak p-value correction). Coefficients and 95% confidence interval of the fitted growth curve, small: *β*_1_=0.69 [0.57,0.79], *β*_2_=0.49 [0.19,0.91], *λ*=0.62 [0.28,1.07]; large: *β*_1_=0.66 [0.59,0.72], *β*_2_=0.94 [0.41,2.03], *λ*=2.48 [1.57,3.32].

**Figure S3:**
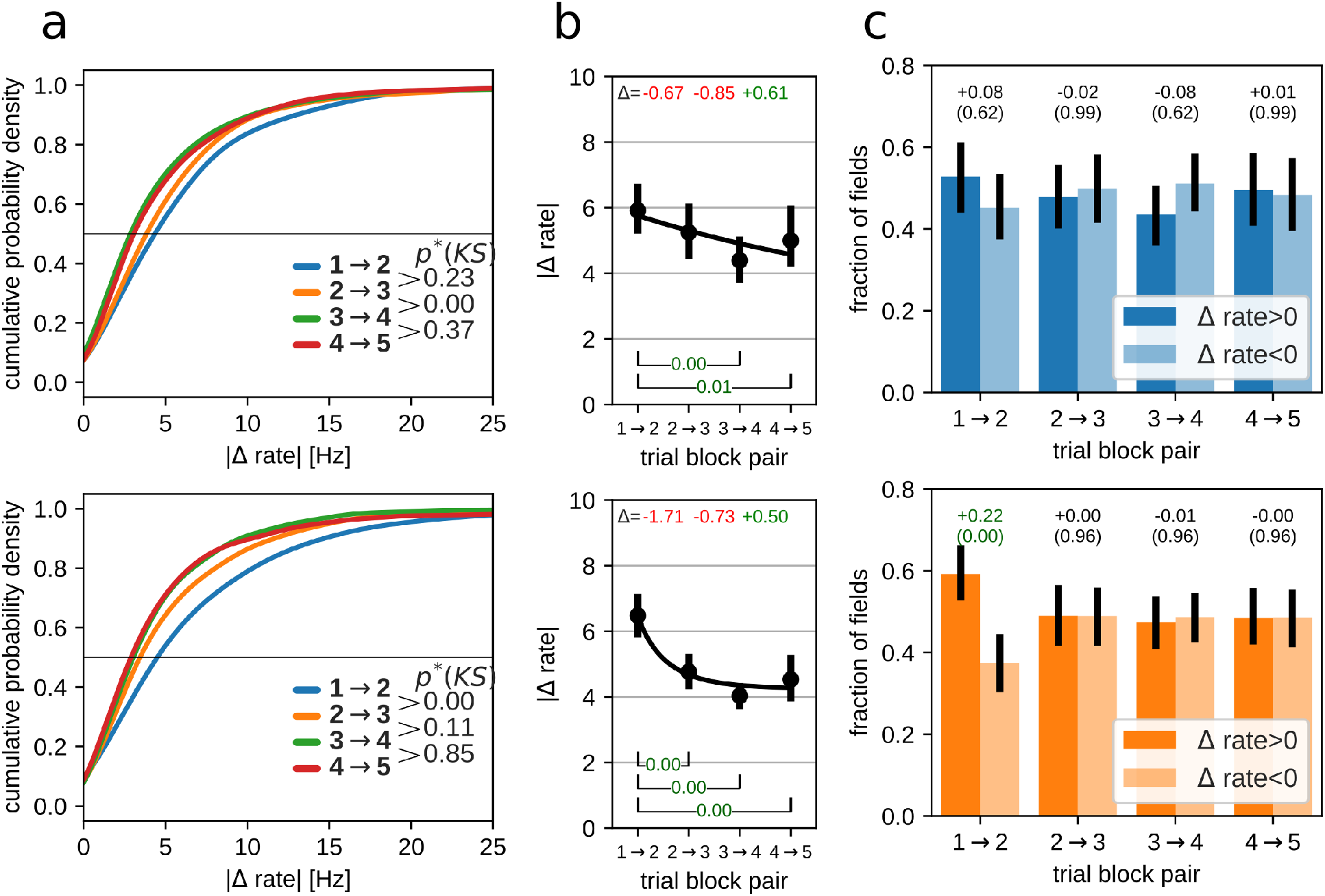
Trial-to-trial changes of in-field firing rate. Same analysis as in figure 4, but excluding emerging and vanishing fields. Top row: small reward, bottom row: large reward. (a) Distributions of the magnitude of in-field rate changes |Δ*rate*| between pairs of trial blocks for all place fields in all sessions. p*(KS) is the Holm-Sidak corrected p-value for Kolmogorov-Smirnov test for equal distributions. (b) Average per-session magnitude of in-field rate changes for pairs of trial blocks. Error bars indicate 95% bootstrapped confidence interval. Text annotations at the top indicate the difference between consecutive trial block pairs. For both reward conditions, Friedman test indicated significant differences between trial-block pairs (small: statistic=14.97, p=0.0018; large: statistic=25.83, p=1×10^−5^). Annotations at the bottom of the plots indicate p-value for significant pairwise differences (Conover posthoc test with Holm-Sidak p-value correction). Coefficients and 95% confidence interval of the fitted growth curve, small: *β*_1_=0.22 [0.16,4.67], *β*_2_=−0.96 [−0.97,−0.24], *λ*=0.00 [0.00,1.26]; large: *β*_1_=4.25 [2.95,4.70], *β*_2_=−0.34 [−0.54,−0.25], *λ*=1.32 [0.29,2.87]. (c) Average fraction of fields with increasing (**dark**) or decreasing (**light**) in-field firing rate. Error bars indicate 95% bootstrapped confidence interval. Text annotations: top row, fraction difference between fields with increasing and decreasing in-field firing rate; bottom row: Holm-Sidak corrected p-value for Wilcoxon signed-rank test between increasing and decreasing rate fractions.

### Immobility across trial blocks

**Figure S4:**
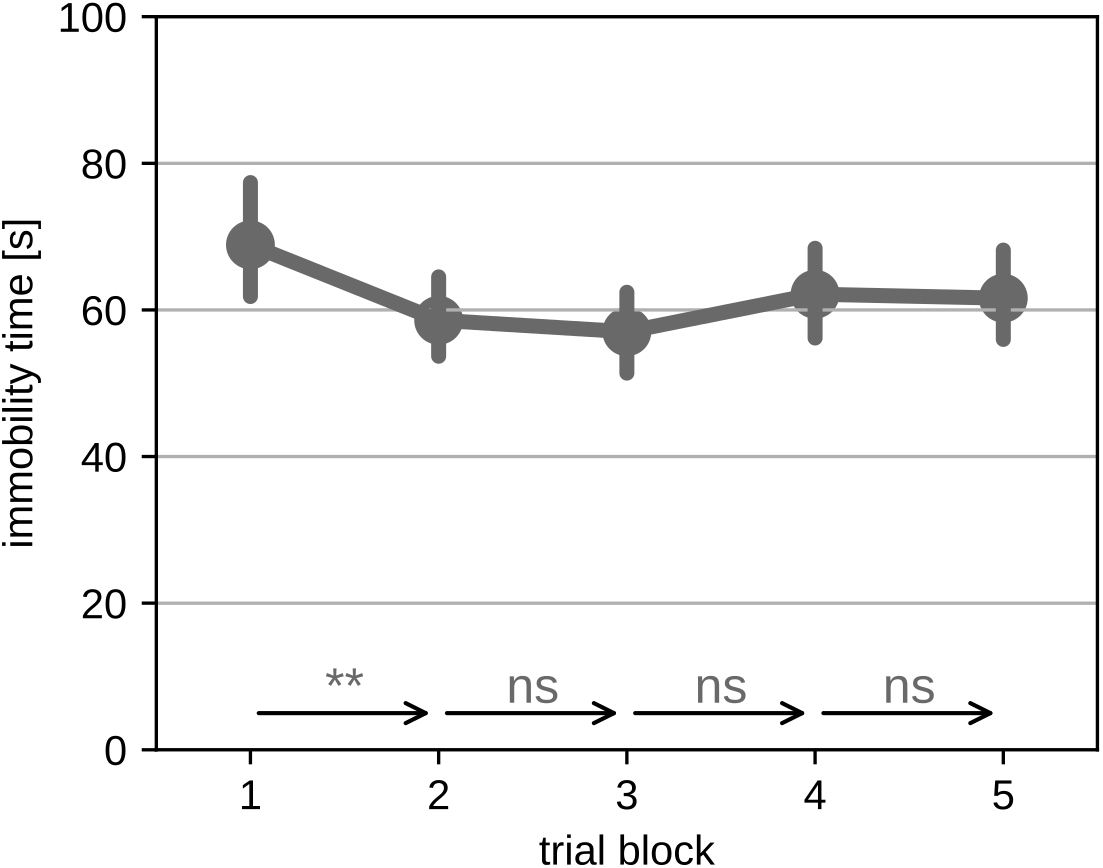
The time spent in a state of immobility mostly remained stable over training. Trial-to-trial changes in average time spent immobile (speed < 5 cm/s). Error bars indicate **95%** bootstrapped confidence interval. Annotations: arrows represent the trial-to-trial transitions annotated with the significance level (Wilcoxon signed-rank test followed by Holm-Sidak correction for multiple tests). **: p<0.01, ns: not significant.

### Replay dynamics across trial blocks: single-trial models

**Figure S5:**
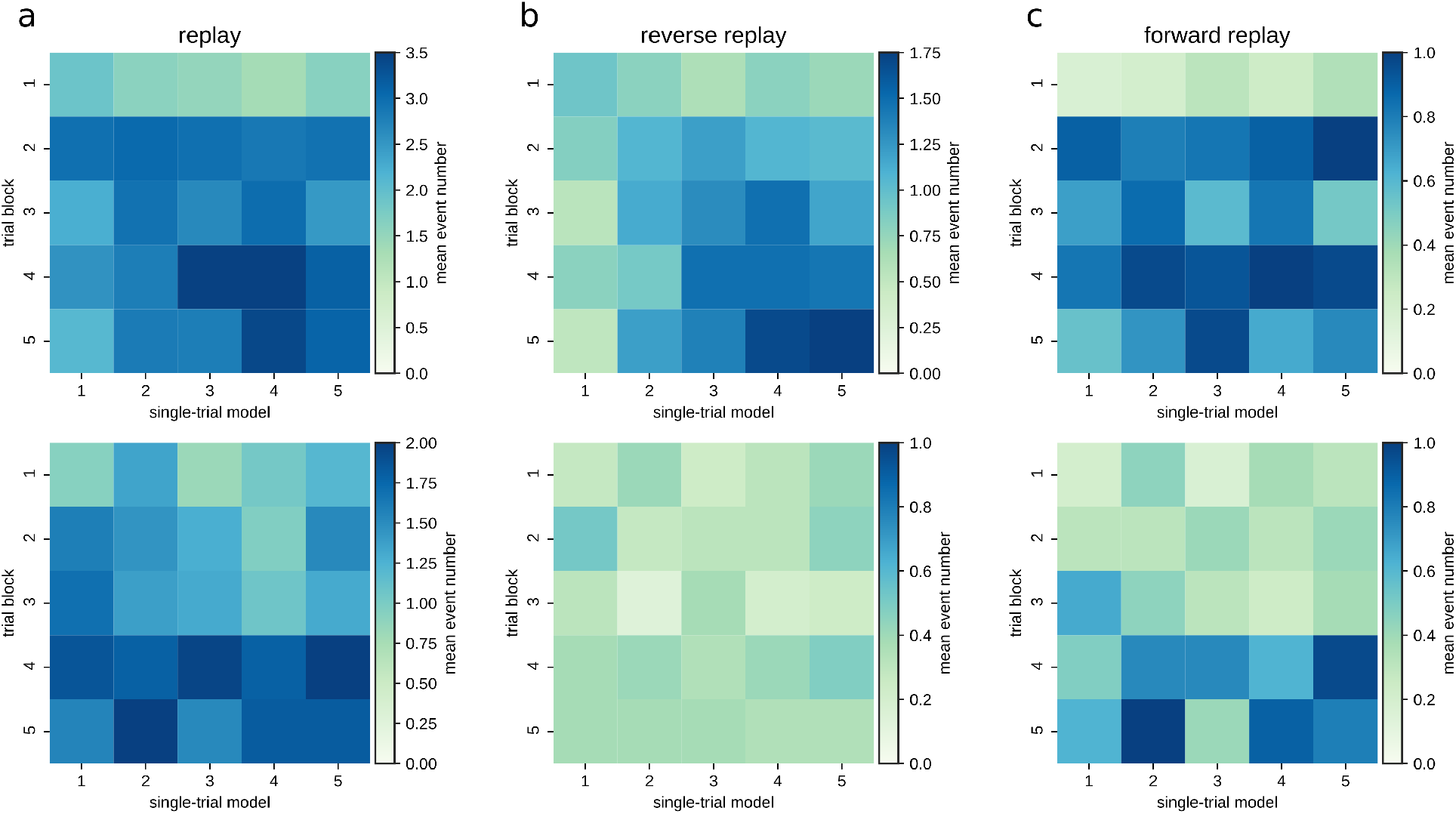
Average number of detected replay events in each trial block for all single-trial encoding models. For all panels, top: replay of large reward environment, bottom: replay of small reward environment. (a) All trajectory replay. (b) Reverse replay. (c) Forward replay.

### Replay dynamics across trial blocks: trial-average model

**Figure S6:**
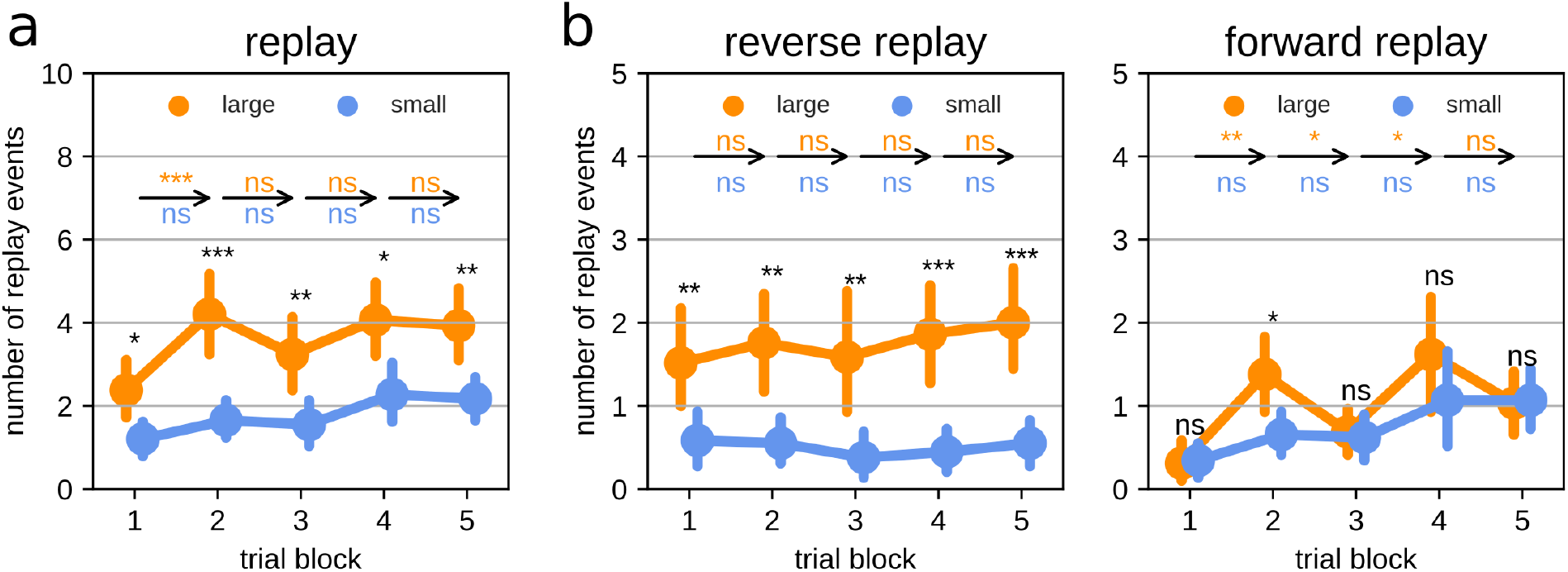
Changes in replay over training and between reward conditions. Replay is decoded using the trial-average model. (a) Number of 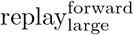 and 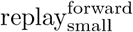 events across trial blocks. (b) Number of reverse (left) and forward (right) replay events across trial blocks. Error bars indicate **95%** bootstrapped confidence interval. Text annotations: arrows represent the trial-to-trial transitions annotated with the significance level separately for the two reward groups. For each trial block, significance level for the difference between the two reward groups is indicated above the data points. All pairwise comparisons were performed with Wilcoxon signed-rank test followed by Holm-Sidak correction for multiple tests. ***: p<0.001; **: p<0.01; *: p<0.05, ns: not significant.

## Notes

### Competing Interest Statement

The authors have declared no competing interest.

